# No compelling evidence that more physically attractive young adult women have higher estradiol or progesterone

**DOI:** 10.1101/136515

**Authors:** Benedict C Jones, Amanda C Hahn, Claire I Fisher, Hongyi Wang, Michal Kandrik, Junpeng Lao, Chengyang Han, Anthony J Lee, Iris J Holzleitner, Lisa M DeBruine

## Abstract

Putative associations between sex hormones and attractive physical characteristics in women are central to many theories of human physical attractiveness and mate choice. Although such theories have become very influential, evidence that physically attractive and unattractive women have different hormonal profiles is equivocal. Consequently, we investigated hypothesized relationships between salivary estradiol and progesterone and two aspects of women’s physical attractiveness that are commonly assumed to be correlated with levels of these hormones: facial attractiveness (N=249) and waist-to-hip ratio (N=247). Our analyses revealed no compelling evidence that women with more attractive faces or lower (i.e., more attractive) waist-to-hip ratios had higher levels of estradiol or progesterone. One analysis did suggest that women with more attractive waist-to-hip ratios had significantly higher progesterone, but the relationship was weak and the relationship not significant in other analyses. These results do not support the influential hypothesis that between-women differences in physical attractiveness are related to estradiol and/or progesterone.

## 1. Introduction

Many researchers have hypothesized that human attractiveness judgments are psychological adaptations for identifying high-quality mates (Grammer et al., 2003; Little et al., 2011; Thornhill & Gangestad, 1999). Researchers have also hypothesized that fertility, as indexed by high levels of estradiol and/or progesterone, is a particularly important aspect of women’s mate quality (Grammer et al., 2003; Little et al., 2011; Thornhill & Gangestad, 1999). Although this proposal has become very influential in the human attractiveness and mate choice literatures, evidence that more physically attractive women have higher estradiol or progesterone is equivocal (Grillot et al., 2014; Jasienska et al., 2004; Law Smith et al., 2006; Puts et al., 2013).

Two studies have investigated putative relationships between women’s facial attractiveness and hormone levels. Law Smith et al. (2006) reported a significant positive correlation between ratings of women’s facial attractiveness and estradiol. They also reported a positive correlation between facial attractiveness and progesterone, although this relationship was not significant. By contrast with Law Smith et al’s results, Puts et al. (2013) found no evidence that women with higher levels of either estradiol or progesterone possessed more attractive faces. To date, evidence that more facially attractive women have higher estradiol or progesterone is therefore inconclusive.

Other studies have tested for evidence that women’s physical attractiveness is positively correlated with estradiol or progesterone by investigating the hormonal correlates of women’s waist-to-hip ratio. Jasienska et al. (2004) reported that women with lower (i.e., more attractive) waist-to-hip ratios had higher estradiol and higher progesterone. However, Grillot et al. (2014) found no evidence for these relationships. To date, evidence that waist-to-hip ratio is associated with sex hormones is therefore also inconclusive.

Given the importance of associations between hormone levels and attractiveness for theories of women’s attractiveness and mate choice, we tested for the hypothesized correlations between salivary estradiol and progesterone and both women’s facial attractiveness and waist-to-hip ratio. Our study is the largest to date to test for putative associations between women’s physical attractiveness and measured hormone levels. Our sample is more than eight times larger than that in Law-Smith et al. (2006) and more than twice as large as that in Jasienska et al. (2004).

## 2. Methods

### 2.1. Participants

We recruited 249 young adult white women for the study (mean age=21.5 years, SD=3.30 years). All participants were students at the University of Glasgow and each completed five weekly test sessions. Participants were recruited only if they were not currently using any hormonal supplements (e.g., oral contraceptives), had not used any form of hormonal supplements in the 90 days prior to their participation, and had never used sunbeds or tanning products. None of the participants reported being pregnant, having been pregnant recently, or breastfeeding. Women participated as part of a larger study on hormonal correlates of women’s behavior (Jones et al., in press a, in press b, in press c).

### 2.2. Face photography and ratings

In each of the five test sessions, each participant first cleaned her face with hypoallergenic face wipes to remove any makeup. Makeup was removed because Law Smith et al. (2006) reported that estradiol and progesterone predicted facial attractiveness in a sample of women not wearing makeup, but not in a sample of women wearing makeup. A full-face digital photograph was taken a minimum of 10 minutes later. Photographs were taken in a small windowless room against a constant background, under standardized diffuse lighting conditions, and participants were instructed to pose with a neutral expression. Camera-to-head distance and camera settings were held constant. Participants wore a white smock covering their clothing when photographed to control for possible effects of reflectance from clothing. Photographs were taken using a Nikon D300S digital camera and a GretagMacbeth 24-square ColorChecker chart was included in each image for use in color calibration.

Following Jones et al. (2015), face images were color calibrated using a least-squares transform from an 11-expression polynomial expansion developed to standardize color information across images (Hong et al., 2001). Note that color calibration of face images eliminates differences across images due to subtle variation in factors such as lighting. It does not reduce differences among images in other aspects of facial coloration. For example, even subtle hormone-linked differences in facial coloration can be measured in images calibrated in this way (Jones et al., 2015). Each image was standardized on pupil positions and masked so that hairstyle and clothing were not visible. The 1245 face images (five images for each of the 249 women) were then rated for attractiveness using a 1 (much less attractive than average) to 7 (much more attractive than average) scale by 14 men and 14 women. Inter-rater agreement for these ratings was high (Cronbach’s alpha=.93). Trial order was fully randomized. The screen was calibrated using an xRite i1 Display Pro colorimeter prior to testing. Simulations (see DeBruine & Jones, 2018) sampling from a population of 2513 raters, each of whom had rated the attractiveness of 102 faces, indicate that >99% of 1000 random samples of 15 raters produced Cronbach’s alphas >.8, indicating high reliability of ratings (90% of all alphas were >.85). Furthermore, increasing the number of raters providing attractiveness ratings has a negligible effect on the mean attractiveness ratings once ratings have been collected from 28 raters (Hehman et al., 2018).

### 2.3. Hormone assays

Participants provided a saliva sample via passive drool (Papacosta & Nassis, 2011) in each test session. Participants were instructed to avoid consuming alcohol and coffee in the 12 hours prior to participation and avoid eating, smoking, drinking, chewing gum, or brushing their teeth in the 60 minutes prior to participation. Saliva samples were frozen immediately and stored at - 32°C until being shipped, on dry ice, to the Salimetrics Lab (Suffolk, UK) for analysis, where they were assayed using the Salivary 17ß-Estradiol Enzyme Immunoassay Kit 1-3702 (M=3.42 pg/mL, SD=1.33 pg/mL; intra-assay CV=7.13%; inter-assay CV=7.45%) and Salivary Progesterone Enzyme Immunoassay Kit 1-1502 (M=143.90 pg/mL, SD=93.33 pg/mL; intra-assay CV=6.2%; inter-assay CV=7.55%). Hormone levels more than three standard deviations from the sample mean for that hormone or where Salimetrics indicated levels were outside the assay sensitivity range were excluded from the dataset (~1.5% of hormone measures were excluded). Reliability of hormone levels across test sessions was good for both estradiol (Cronbach’s alpha=.90; Intraclass correlation coefficient=.46) and progesterone (Cronbach’s alpha=.91; Intraclass correlation coefficient=.58).

### 2.4. Body measures

In one of the five test sessions, waist and hip circumferences were measured from 247 of the women by one researcher. Two women chose not to have waist and him circumferences measured. Waist and hip circumferences were used to calculate waist-to-hip ratio (M=0.75, SD=0.05).

## 3. Results

A linear mixed model was used to investigate the relationship between facial attractiveness and hormone levels. Analyses were conducted using R version 3.3.2 (R Core Team, 2016), with lme4 version 1.1-13 (Bates et al., 2014) and lmerTest version 2.0-33 (Kuznetsova et al., 2013). To create mean (i.e., trait) hormone values for our analyses, hormone levels were averaged across test sessions for each woman, centered on the grand mean, and scaled so the majority of the distribution for each hormone varied from −.5 to .5 (this was done by dividing values by a constant, is done simply to facilitate calculations in the linear mixed models, and has no material effect on the results). To create current (i.e., state) hormone values for our analyses, values for each hormone were centered on their subject-specific means and scaled using the same scaling constants as above. The linear mixed model predicted face image ratings with current (i.e., state) estradiol, current (i.e., state) progesterone, rater sex (effected coded so that +0.5 was male and −0.5 was female), and their interactions entered as predictors. Mean (i.e., trait) estradiol, mean (i.e., trait) progesterone, rater sex, and their interactions were also entered as predictors. Interactions between estradiol and progesterone were included following Puts et al. (2013). Random intercepts were specified for rater, stimulus woman (i.e., each woman whose face images were used as stimuli), and individual face image. Random slopes were specified maximally, following Barr et al. (2013) and Barr (2013). The model is fully described in our supplemental materials, along with results of simplified models testing for effects of current and mean hormone levels separately (see https://osf.io/qd9bv/). Data are also available at https://osf.io/qd9bv/. Full results are shown in Table 1.

**Table 1.** Results of linear mixed model testing for within-woman and between-women hormone-attractiveness correlations.

|  | estimate | 95% CI | SE | t | p |
| --- | --- | --- | --- | --- | --- |
| current estradiol | -0.01 | -0.10, 0.08 | 0.05 | -0.23 | .82 |
| current progesterone | -0.07 | -0.14, 0.01 | 0.04 | -1.66 | .10 |
| rater sex | -0.70 | -1.20, -0.20 | 0.26 | -2.73 | .01 |
| mean estradiol | -0.15 | -0.56, 0.26 | 0.21 | -0.73 | .47 |
| mean progesterone | -0.06 | -0.63, 0.51 | 0.29 | -0.21 | .83 |
| current estradiol x current progesterone | -0.54 | -1.01, -0.06 | 0.24 | -2.20 | .03 |
| current estradiol x rater sex | 0.02 | -0.13, 0.17 | 0.08 | 0.23 | .82 |
| current progesterone x rater sex | -0.02 | -0.16, 0.11 | 0.07 | -0.36 | .72 |
| mean estradiol x mean progesterone | 0.69 | -1.70, 3.08 | 1.22 | 0.56 | .57 |
| mean estradiol x rater sex | -0.00 | -0.11, 0.11 | 0.05 | -0.01 | .99 |
| mean progesterone x rater sex | -0.09 | -0.24, 0.06 | 0.08 | -1.17 | .24 |
| current estradiol x current progesterone x rater sex | 0.42 | -0.45, 1.28 | 0.44 | 0.94 | .36 |
| mean estradiol x mean progesterone x rater sex | 0.23 | -0.65, 1.11 | 0.45 | 0.52 | .61 |

No between-women hormone-attractiveness correlations were significant. However, there was a significant interaction between the effects of current estradiol and current progesterone (estimate=−0.54, 95% CI=−1.01, −0.06, SE=0.24, t=−2.20, p=.030). Although weak, this interaction indicated that within-woman attractiveness was particularly high both when current estradiol was high and current progesterone was simultaneously low and when current estradiol was low and current progesterone was simultaneously high (see Figure 1).

**Figure 1.**
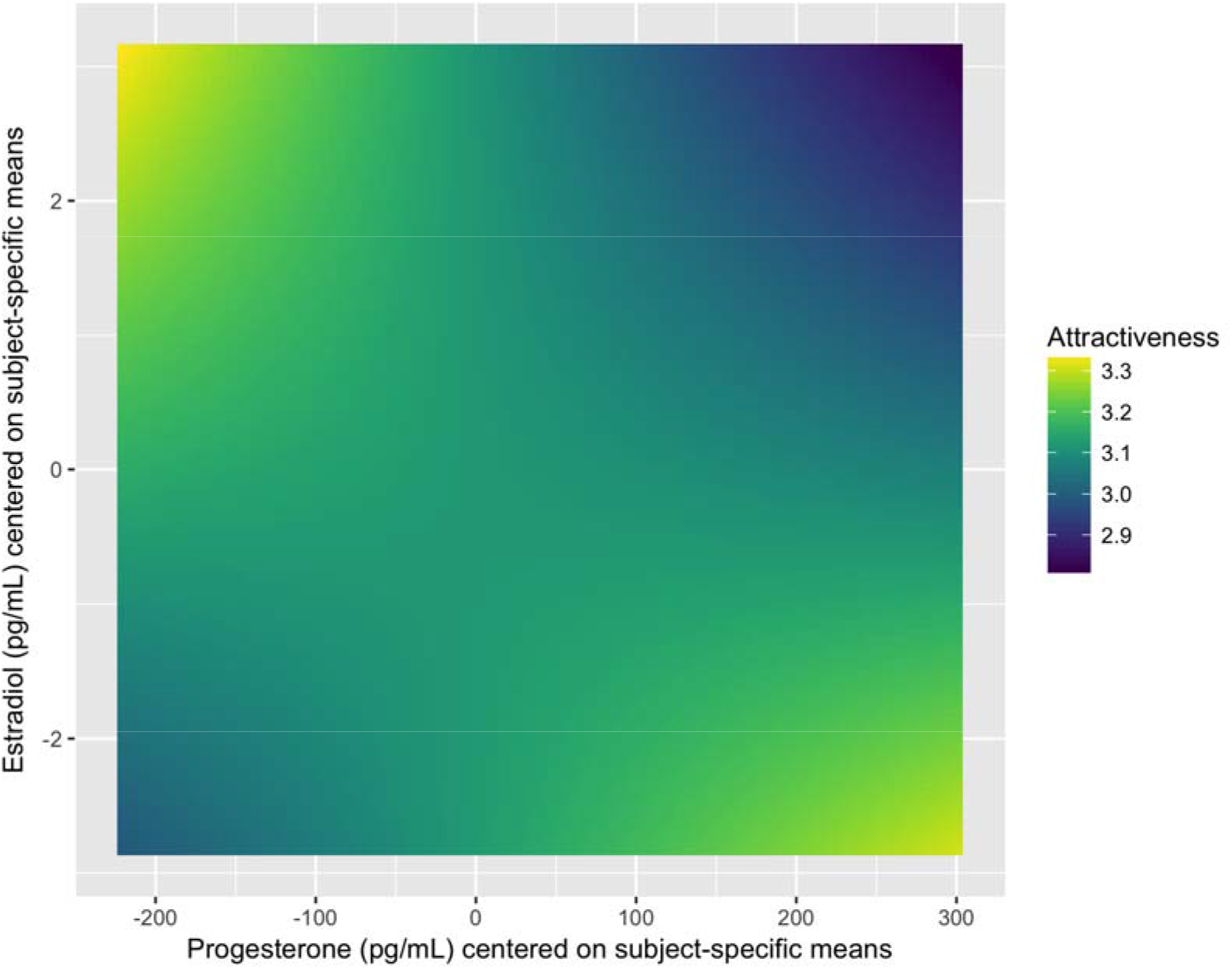
The interaction between current estradiol and current progesterone. The heat map shows predicted values for attractiveness based on the model tested and for the range of estradiol and progesterone values in our dataset.

Since we had only one waist-to-hip ratio measure for each woman, we simply tested for significant correlations between waist-to-hip ratio and both mean estradiol and mean progesterone. There was a significant positive correlation between waist-to-hip ratio and mean estradiol (r=.23, 95% CI=.11, .34, N=247, p<.001). The correlation between waist-to-hip ratio and mean progesterone was not significant (r=−.07, 95% CI=−.20, 0.05, N=247, p=.24).

Next, we repeated the between-women analyses of facial attractiveness and waist-to-hip ratio, this time controlling for between-women differences in body mass index (BMI). Although we observed between-women hormone-BMI correlations, controlling for BMI did not alter the patterns of results described above. These analyses are described in full in our supplemental materials (https://osf.io/qd9bv/). One woman chose not to have her height and weight measurements taken so her data could not be included in these analyses.

In addition to the analyses described above, we investigated the relationships between attractiveness and average hormone levels and between WHR and average hormone levels using a Bayesian analysis with a multivariate latent model. Details of this analysis and full results are reported at https://osf.io/qd9bv/. The repeated measurement for each female subject (estradiol and progesterone) and the repeated measurement from the raters were modeled as the realization of the unobserved latent variables with a mixed effect formulation. Similar to the results above, the correlation between waist-to-hip ratio and the latent estradiol level was estimated at r=.261 [.138, .386] (bracket shows 95% highest posterior density interval). The correlation between waist-to-hip ratio and the latent progesterone was r=−.022 [−.160, .113]. The correlation between the latent estradiol level and latent attractiveness rating was r=−.044 [−.182, .087]. The correlation between the latent progesterone level and latent attractiveness rating was r=−.077 [−.215, .058].

In response to a reviewer’s suggestions, we tested for correlations between hormone levels and both attractiveness and waist-to-hip ratio using each participant’s maximum progesterone level and maximum estradiol level. We did this to address concerns about the extent to which average hormone levels might be biased by some participants being tested more often in particular cycle phases. These analyses showed no significant relationships between maximum hormone levels and facial attractiveness (both absolute r<.03, both p>.64). Consistent with our previous analyses, women with higher maximum estradiol had significantly higher waist-to-hip ratios (r=.16, p<.010). Although the relationship was weak, women with higher maximum progesterone had significantly lower waist-to-hip ratios (r=−.14, p=.020). However, restricting the data set to women with maximum progesterone levels greater than 250 pg/mL (i.e., those showing evidence of having ovulated prior to their maximum progesterone level being measured) altered the pattern of results for waist-to-hip ratio, however. In these analyses, women with higher maximum estradiol still had significantly higher waist-to-hip ratios (r=.31, p<.001), but the relationship between maximum progesterone waist-to-hip ratio was now not significant (r=−.05, p=.63). That the correlation between progesterone and waist-to-hip ratio was only significant in one of our analyses suggests it is not robust.

## 4. Discussion

Here we investigated possible relationships between salivary estradiol and progesterone and both women’s facial attractiveness and waist-to-hip ratio. We carried out these analyses to test the influential hypothesis that more physically attractive women have higher estradiol and progesterone (Jasienska et al., 2005; Law Smith et al., 2006). We found no compelling evidence that women with higher facial attractiveness or lower (i.e., more attractive) waist-to-hip ratios had higher levels of estradiol or progesterone^1^. In fact, we actually found that women with higher (i.e., relatively unattractive) waist-to-hip ratios had higher levels of estradiol. Thus, our results do not replicate those of previous studies with smaller sample sizes reporting significant correlations between hormone levels and either facial attractiveness (Law Smith et al., 2006) or waist-to-hip ratio (Jasienska et al., 2004). Our results do not then support the influential hypothesis that between-woman differences in physical attractiveness are correlated with estradiol and/or progesterone. The Bayesian analyses we carried out also supported this conclusion.

We observed no evidence that more attractive women had higher estradiol or progesterone levels. However, our analysis of facial attractiveness ratings suggested that within-woman changes in facial attractiveness were associated with within-woman changes in hormone levels. Women’s facial attractiveness subtly increased both when current estradiol was high and current progesterone was simultaneously low and when current estradiol was low and current progesterone was simultaneously high. This result partially replicates Puts et al. (2013), who found that attractiveness was increased when current estradiol was high and current progesterone was simultaneously low.

The combination of high estradiol and low progesterone is characteristic of the fertile phase of the menstrual cycle (Gangestad & Haselton, 2015). Consequently, Puts et al. (2013) proposed that the increased attractiveness that they observed when women were in this hormonal state supported the hypothesis that women’s attractiveness subtly increases during the fertile phase of the menstrual cycle. However, Puts et al. (2013) compared attractiveness during the late follicular and mid-luteal phases of the menstrual cycle only. Because relatively high levels of both progesterone and estradiol characterize the mid-luteal phase of the menstrual cycle, Puts et al. (2013) are unlikely to have sampled women when estradiol was low and progesterone was simultaneously high. By contrast, we sampled women at weekly intervals over an entire menstrual cycle, allowing us to capture a greater range of hormonal states. Importantly, our results showing that attractiveness increased both when current estradiol was high and current progesterone was simultaneously low and when current progesterone was high and current estradiol was simultaneously low suggest that hormone-linked increases in facial attractiveness are not necessarily unique to hormonal states associated with high fertility. Thus, our results for within-woman hormone-attractiveness correlations do not necessarily support the hypothesis that hormone-linked within-woman changes in attractiveness are fertility signals or the hypothesis that they are imperfectly concealed cues of ovulatory status (see Haselton & Gildersleeve, 2016 for a discussion of these two hypotheses). Although we acknowledge that the combination of high levels of progesterone and low levels of estradiol is a relatively unusual combination of hormone levels in textbook plots of hormonal changes over the menstrual cycle, graphing the distribution of these hormones in our dataset suggests it is not particularly uncommon in our sample. This is consistent with other recent work suggesting that only ~38% of young adult women show textbook hormonal profiles (see Marcinkowska et al., 2018).

Havlicek et al. (2015a, 2015b) posited that within-woman hormone-attractiveness correlations might simply be functionless byproducts of between-women hormone-attractiveness correlations. We agree with Havlicek et al. that the within-subject changes in attractiveness observed in the current study (and in other studies reporting changes in facial attractiveness during the menstrual cycle) may be too subtle to directly influence men’s mate choices. Nonetheless, that we observed significant within-woman, but not between-women, hormone-attractiveness correlations does not support Havlicek et al’s hypothesis that within-woman hormone-attractiveness correlations are functionless byproducts of between-women hormone-attractiveness correlations. We note here that, while we suggest that within-woman hormone-attractiveness correlations are unlikely to simply be functionless byproducts of between-women hormone-attractiveness correlations, we remain open to the possibility that they are functionless byproducts of within-woman hormonal changes, rather than fertility signals.

Could differences in methodologies across studies explain the inconsistent results for hormone-attractiveness correlations? For example, Law-Smith et al. (2006) and Jasienska et al. (2004) both controlled for effects of cycle-linked within-woman variation in hormone levels when testing for between-women hormone-attractiveness correlations. Law-Smith et al. (2006) did this by measuring estradiol and progesterone from the late follicular and luteal menstrual cycle phases, respectively. Jasienska et al. (2004) did this by averaging hormone levels measured daily and also by considering luteal-phase progesterone levels only. Thus, the differences between our findings and their results could be due to the possible effects of cycle phase on between-women hormone differences being less well controlled in our study than in Jasienska et al. (2004) and Law-Smith et al. (2006). Although the logic of this explanation is appealing, we think this explanation is unlikely, Our results are consistent with those of other studies that controlled for effects of cycle-linked changes in hormone levels when investigating the between-women hormone-attractiveness correlations, but also found no compelling evidence that more attractive women had higher levels of estradiol or progesterone (Grillot et al., 2014; Puts et al., 2013).

The positive correlation between estradiol and waist-to-hip ratio that we observed in the current study is surprising. However, we note that the pattern of results across studies on this issue (i.e., the negative correlation reported by Jasienska et al., the null result reported by Grillot et al., and the positive correlation observed in the current study) is arguably what one would expect if there are no reliable correlations between waist-to-hip ratio and these sex hormones (i.e., we suggest that both the positive correlation in our study and the negative correlation reported by Jasienska et al. may both be false positives). We also suggest that it is unlikely that our non-replication of Jasienska et al’s results for waist-to-hip ratio are a consequence of having only a single measurement of waist-to-hip ratio taken by an individual researcher for each woman. While taking more measurements could reduce measurement error, it is unlikely that this source of error is causing the strikingly different patterns of results seen in Jasienska et al. and the current study.

In conclusion, our analyses provide no compelling evidence that women with more attractive faces or waist-to-hip ratios have higher estradiol or progesterone. Importantly, these null results do not support the popular and influential hypothesis that women’s physical attractiveness is a marker for their estradiol and/or progesterone levels, at least among young adult women.

## 5. Acknowledgments

We thank Sean Murphy, Elizabeth Necka, Craig Roberts, and Jan Havlicek for constructive comments and discussion.

## Footnotes

1 Although one analysis showed a significant negative correlation between maximum progesterone and waist-to-hip ratio, the correlation was weak and not evident in our other analyses. This pattern of results suggests the relationship between progesterone and waist-to-hip ratio is not robust and may be a false positive,

